# The dynamics of prion spreading is governed by the interplay between the non-linearities of tissue response and replication kinetics

**DOI:** 10.1101/2024.05.01.592001

**Authors:** Basile Fornara, Angélique Igel, Vincent Béringue, Davy Martin, Pierre Sibille, Laurent Pujo-Menjouet, Human Rezaei

**Author notes:** Corresponding authors and senior authorships: Laurent Pujo-Menjouet, Human Rezaei, MAP2, VIM, INRAe, Domaine de Vilvert, 78352, Jouy-en-Josas, France.

## Abstract

Prion diseases, or Transmissible Spongiform Encephalopathies (TSE), are neurodegenerative disorders caused by the accumulation of misfolded conformers (PrP^Sc^) of the cellular prion protein (PrP^C^). During the pathogenesis, the PrP^Sc^ seeds disseminate in the central nervous system and convert PrP^C^ leading to the formation of insoluble assemblies. As for conventional infectious diseases, variations in the clinical manifestation define a specific prion strain which correspond to different PrP^Sc^ structures. In this work, we implemented the recent developments on PrP^Sc^ structural diversity and tissue response to prion replication into a stochastic reaction-diffusion model using an application of the Gillespie Algorithm. We showed that this combination of non-linearities can lead prion propagation to behave as a complex system, providing an alternative to the current paradigm to explain strain specific phenotypes, tissue tropisms and strain co-propagation while also clarifying the role of the connectome in the neuro-invasion process.

## Introduction

Prion diseases, or Transmissible Spongiform Encephalopathies (TSE), are a class of untreatable fatal neurodegenerative disorders caused by the accumulation and aggregation in the central nervous system of misfolded conformers (PrP^Sc^) of the host-encoded membrane-anchored prion protein (PrP^C^). Throughout the course of the infection, the PrP^Sc^ assemblies replicate by converting PrP^C^ into PrP^Sc^, inducing the extracellular accumulation of the infectious conformers leading to the formation of pathogenic aggregates ^1,2^ . This autocatalytic conversion of PrP^C^ into PrP^Sc^ constitutes the basis of the prion paradigm which has been extended to Parkinson’s and Alzheimer’s diseases to describe the dissemination process and the progression of these pathologies ^3^.

As for conventional infectious diseases in which variations in the clinical manifestation define the pathogenic strain, in prion diseases, variations in incubation period, pattern of neural loss, PrP^Sc^ deposition and biochemical properties of PrP^Sc^ assemblies define a specific prion strain ^4,5^. The biochemical properties include the type of fragments generated after proteolysis ^6,7^, apparent resistance to unfolding ^8,9^ and size distribution of PrP^Sc^ assemblies at the terminal stage of the disease ^10–12^. While the differences in physicochemical properties can be explained by the variations in the structure of PrP^Sc^ which have recently been identified ^13,14^, the link between structure and clinical manifestation has yet to be uncovered.

The spatiotemporal progressions of tauopathy and misfolded protein deposition in Alzheimer’s and Parkinson’s diseases follow the neuronal connectome in a well-established manner called Braak’s staging ^15,16^. In prion diseases however, the contribution of the connectome to the disease progression and histological patterning is far to be valid ^17^. Indeed, neither the pattern of PrP^Sc^ deposition nor the lesional profile follow the connectome as they appear to depend solely on the biochemical properties of the strain ^17^. Based on experimental observations, a correlation between levels of PrP^C^ expression and PrP^Sc^ deposition is far to be a general rule ^18,19^. To explain how the strain properties define the lesional profile and PrP^Sc^ deposition, a prevalent hypothesis lacking any experimental basis relies on the tropism between a given strain and cells expressing a specific conformer of PrP^C^ (local PrP^C^ conformome hypothesis) ^20^. In this hypothesis, the physicochemical properties of PrP^C^ are cell-dependent ^21^ and prion strains would therefore favor specific brain regions. However, several experimental observations dispel this theory. The first one is the absence of PrP^C^ folding intermediates and the highly reversible two-step unfolding/refolding process of the native state of PrP ^22,23^ which goes against the existence of PrP^C^ conformers. The second one is the high efficient rate of conversion in Protein Misfolding Cyclic Amplification (PMCA) conditions with distinct cell lysates coming from different brain regions ^24^ refuting cell-specific templating differences. The third one is the dependency of PrP^Sc^ deposition pattern on the inoculation pathway and inoculated dose which also allows us to discard the potential impact of local cofactors ^25–27^. The fourth one is the spatial and temporal invariance of the seeding activity during the progression of the disease indicating that every brain area presents quasi-similar replication efficiency for a given strain ^28^.

Once the local PrP^C^ conformome hypothesis is discarded, another explanation to strain-specific patterns relies on the kinetics at the base of the replication and the dissemination. It was proven both mathematically and biochemically that a reaction-diffusion system where at least two diffusible reactants interact with each other through non-linear feedback (e.g., catalysis) is able to self-organize in defined patterns ^29,30^. Recent studies on the early stage of prion replication revealed the formation of two structurally distinct sets of PrP^Sc^ assemblies, chemically tied by a secondary templating pathway ^31^. The study of the dynamics of infectious recombinant prions highlighted catalytic conformational exchanges between different PrP^Sc^ subpopulations ^32^. This particular dynamicity of different PrP^Sc^ subpopulations combined with a local tissue response affecting PrP^C^ synthesis, such as the Unfolded Protein Response (UPR) ^33–37^ or variable PrP^C^ production rates ^19^, could constitute a new and plausible hypothesis to explain prion strain phenotypes or how two different prion strains with different rates of replication could co-propagate within the same individual and escape best replicator selection ^21,38–40^. Indeed, co-propagation is often observed in natural prion infections or when adapting a strain to a new host ^40–42^. During co-propagation, the two strains involved can influence one another, typically occurring between a fast strain (with a short incubation time) and a slow strain (with a long incubation time). The interference can cause a prolongation of the incubation period or even a blockage of the fast strain^43,44^. While it is widely believed that this interference is primarily driven by hypothetical cofactors or competition between the co-infecting strains for PrP^C^, the exact molecular mechanisms remain unclear. Therefore, in the present work, we implemented the recent developments on PrP^Sc^ dynamicity and tissue response to prion replication into a stochastic reaction-diffusion model to study the neuro-invasion process. We showed that this combination provides a valid alternative to explain prion strain phenotypes, copropagation and tissue tropisms.

## Materials and Methods

### Dynamics of the coexistence of different PrP^Sc^ subpopulation and kinetic scheme of their replication

The kinetic model of prion assemblies used in this work is based on the recent evidence of structural diversification occurring during replication ^31^ and the dynamicity of different PrP^Sc^ subpopulations ^32,45,46^. Regardless of the strain, prion replication generates two structurally distinct sets of assemblies called PrP^Sc^A and PrP^Sc^B_i_ (from now on called respectively *A* and *B_i_*). The *A* and *B_1_* species are both dimers of PrP differing in structure and *B_i_* is a condensate of *B_1_* where i represents the number of *B_1_* elementary units it comprises i.e. its size ^31^. Both subassemblies have the ability to spread the strain information from which they are issued. While *B_i_* grows by one unit during replication, *A* replicates by creating a copy of itself, its size remaining constant ^31^. As we consider that *B_i_* assemblies replicate at their extremities, the number of templating interfaces remains the same regardless of their size, therefore the templating of *B_i_* was chosen to be independent of size. Additionally, bioassay experiments proved that the specific infectivity (i.e. replication rate) of subpopulation *A* is substantially higher than that of *B_i_* ^31^.

Biochemical characterization of different prion strains established the existence of a constitutional equilibrium between *B_i≥2_* and *B_1_* assemblies ^45,46^. Based on oscillatory behavior observed during relaxation kinetics experiments, Mezache and colleagues completed this model with autocatalytic processes between *A* and *B_i_* subpopulations ^32^. The complete kinetic model is reported in Fig.1a.

**Fig.1.**
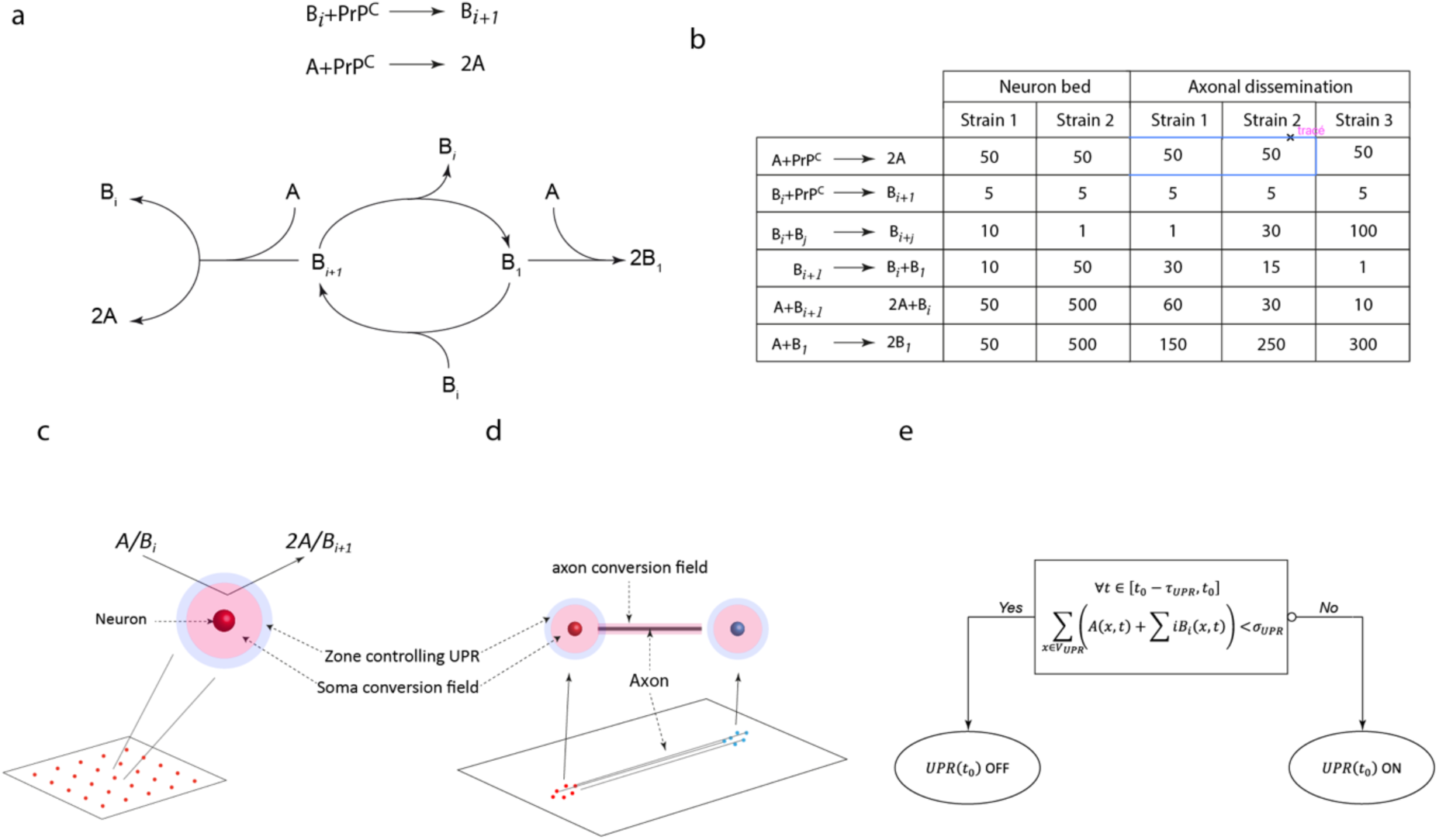
Simulation configurations: (a) Kinetic scheme describing prion replication and structural diversification. The replication process, independently of the strain, generates two conformationally distinct types of PrP^Sc^ assemblies: small oligomeric objects PrP^Sc^A (denoted as *A*) and assemblies capable of condensing PrP^Sc^B_i_ (denoted as *B_i_*) ^31^, *i* referring to the size of the object. These two subpopulations undergo catalytical exchanges according to the kinetic scheme ^32^. Table (b) summarizes the kinetic parameters of modeled strains used in our two types of spreading simulations: on a homogeneous neuron bed (c) and in the axonal dissemination between two clusters (d). The surrounding area of a neuron consists of two zones. The replication field (in pink) defines the zone where replication of *A* and *B_i_* subpopulations can occur, it is located around the somas and axons. The UPR controlling area (in blue) defines the zone around the soma where if the number of assemblies (*A* + ∑ *iB_i_*) exceeds a threshold, denoted *σ*, the UPR activates until the number of assemblies has remained under the threshold for a lag duration *τ*. While the UPR is activated, no templating reaction can occur in the associated replication area of the neuron. This behavior is summarized in the UPR functional diagram (e).

### In silico modeling of strain phenomenon

Recent developments in structural biology highlighted clear differences between PrP^Sc^ fibrils from distinct prion strains^13,14^. It is thus expected that PrP^Sc^ assemblies from two prion strains differ in their physico-chemical properties. In our simulations, we defined a prion strain as the set of kinetic parameters governing the intrinsic dynamicity of the assemblies and their replication rate through the previously described kinetic scheme. Therefore, kinetic parameters defining the modeled strains used in this work are reported in Fig.1b.

### Tissue representation, neuron bed, templating field and tissue response to prion replication and accumulation

While PrP^Sc^ assemblies are extracellular, PrP^C^ is mostly located on the surface of cells attached by a glycosylphosphatidylinositol (GPI) anchor ^47^, templating reactions therefore predominantly occur in the vicinity of cells. In our model, replication of both *A* and *B_i_* subassemblies can only happen inside a zone around neurons we call templating field (Fig.1c), inside which we consider PrP^C^ to be in excess and evenly distributed. On the surface of neurons, PrP^C^ is more abundant around the somas than along the axons ^48^. In our model, this difference translated into the templating field being wider around the somas than the axons.

As reported in the literature, prion accumulation in the vicinity of a cell induces the UPR, through either PrP^Sc^ endocytosis pathways ^49^ or extracellular sensors ^33^, which leads to the down regulation of PrP^C^ synthesis in the cell ^33–37^. Due to the UPR activation relying on the endoplasmic reticulum signaling pathways, we assumed that replication and accumulation of prions along the axons do not contribute to the UPR mechanism. We therefore defined a zone surrounding the soma of the neuron where accumulation of misfolded proteins transiently induces UPR activation ^50^ (Fig.1c&d).

Like most cell regulation systems, the UPR is bistable ^50–52^. In our model, it is triggered when the number of misfolded assemblies (*A* + ∑*_i_^n^ B_i_*) in the UPR control zone exceeds a threshold (denoted *σ*) ^52^. While the UPR of a neuron is activated, no templating reactions can occur in its associated templating field. The resilience (re-activation of templating field) occurs when the total number of misfolded assemblies in the control zone remains under the threshold for a delay *τ_R_*(Fig.1d). Both *σ* and *τ_R_* are parameters specific to a given cell-type and could characterize a given brain area.

To simulate the evolution of prion assemblies on a brain area, we used a uniform square grid of neurons sharing the same UPR parameters. The tissue parameters defining a modeled brain area selected in this work were *N_neur_*, the parameter for the number of neurons in the *N_neur_*-by-*N_neur_* grid, and the UPR threshold *σ*. To mimic the dissemination between two brain areas we had two clusters of neurons unilaterally connected by axons (Fig.1d). The tissue parameters studied here were the direction (anterograde or retrograde) and the number of axons linking the two groups.

### Gillespie stochastic reaction-diffusion modeling

To compute the evolution of the reaction-diffusion model, we used an application of the Gillespie algorithm ^53,54^. The space was divided up into voxels where chemical reactions can only happen between two particles within the same voxel and templating reactions inside of an active templating field. Diffusion was added to the algorithm as a jump process from one voxel to a neighbor whose rate depends on the size of the voxels and the diffusion constant of the particle. A detailed tutorial is provided as supporting information (SI1).

In our model, diffusivity of assemblies is inversely proportional to their size (equation 1) and independent of space (both position and direction). This means that when a diffusion event is selected by the algorithm, the particle has equal chances to move to any of the 4 adjacent voxels.

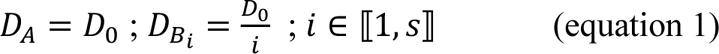

In all our simulations, we chose *D*_0_ = 1000.

The domain in which assemblies evolved was square-shaped and divided in a regular mesh of *div* × *div* square voxels of length *L*. In 2D (4 neighbors), the probability of a particle of diffusivity *D* to exit a voxel of size *L* is 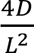, with equal chances of going in any direction. Parameter *L* was chosen to ensure the validity of the Gillespie algorithm which requires that diffusion events greatly outnumber the other reactions.

Assemblies can exit the system by diffusing outside the domain through one of the edges. This is the only way in the model for assemblies to be eliminated and accounts for the spread of elements to other regions as well as clearance of more diffusive objects in vivo. In simulations of brain areas, we kept the distance between the outer layer of the neuron grid and the edges of the domain constant to ensure comparable rates of clearance of assemblies regardless of the neuron density parameter.

### Initial conditions, size of assemblies

While *A* assemblies have a size of 1, *B_i_* can condensate up to a maximum size *s*. This means that the templating and condensing reactions for *B_i_* are constrained:

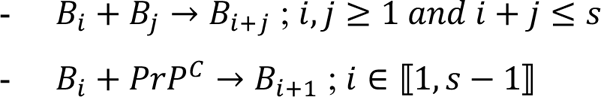

In all our simulations, we chose *s*=10.

All our simulations were seeded with the same quantity and distribution of PrP^Sc^ except for the exploration of initial seed amount where the amount had been increased 100-fold. In brain area simulations, the seed was injected in the center of the neuron grid whereas in dissemination experiments through the neural network, it started at either efferent or afferent groups of neurons.

## Results

### Interplay between local tissue properties and PrP^Sc^ assemblies’ dynamicity

To determine how prion dynamicity and local tissue response are intricated, we computed the evolution of PrP^Sc^ assemblies ruled by the kinetic scheme detailed in the method on a homogeneous bed of neurons where the expression of PrP^C^ was modulated by the UPR. We chose two sets of strain parameters and two sets of tissue parameters (Fig.1b) and then compared the behaviors of the different strain-tissue combinations. While strains 1 and 2 share the same templating activities, due to their intrinsic dynamicity, the balance of subpopulation *B_i_* in strain 1 is shifted more towards the larger assemblies compared to strain 2. Tissues 1 and 2 differ in their density of neurons and UPR threshold: tissue 1 is a 3×3 neuron grid with a low UPR threshold (*σ*_1_ = 100) and tissue 2 is a 5×5 neuron grid with a higher UPR threshold (*σ*_2_ = 300).

Computing the evolution of strain 1 on tissue 1 (*S1T1*) revealed the systematical elimination of subpopulation *B_i_* after a short transient phase while the remaining amount of *A* assemblies adopted an oscillatory behavior (Fig.2a and SI2). The tissue response induced by the sustained evolution of strain 1 displayed periodic synchronized UPR pulses of all the neurons but we noticed with the help of Fig.3b that neurons closer to the middle spent more time activated on average. The reproducibility of these observations over fifty independent simulations suggested the convergence of the evolution of *S1T1* and the stability of this equilibrium.

**Fig.2.**
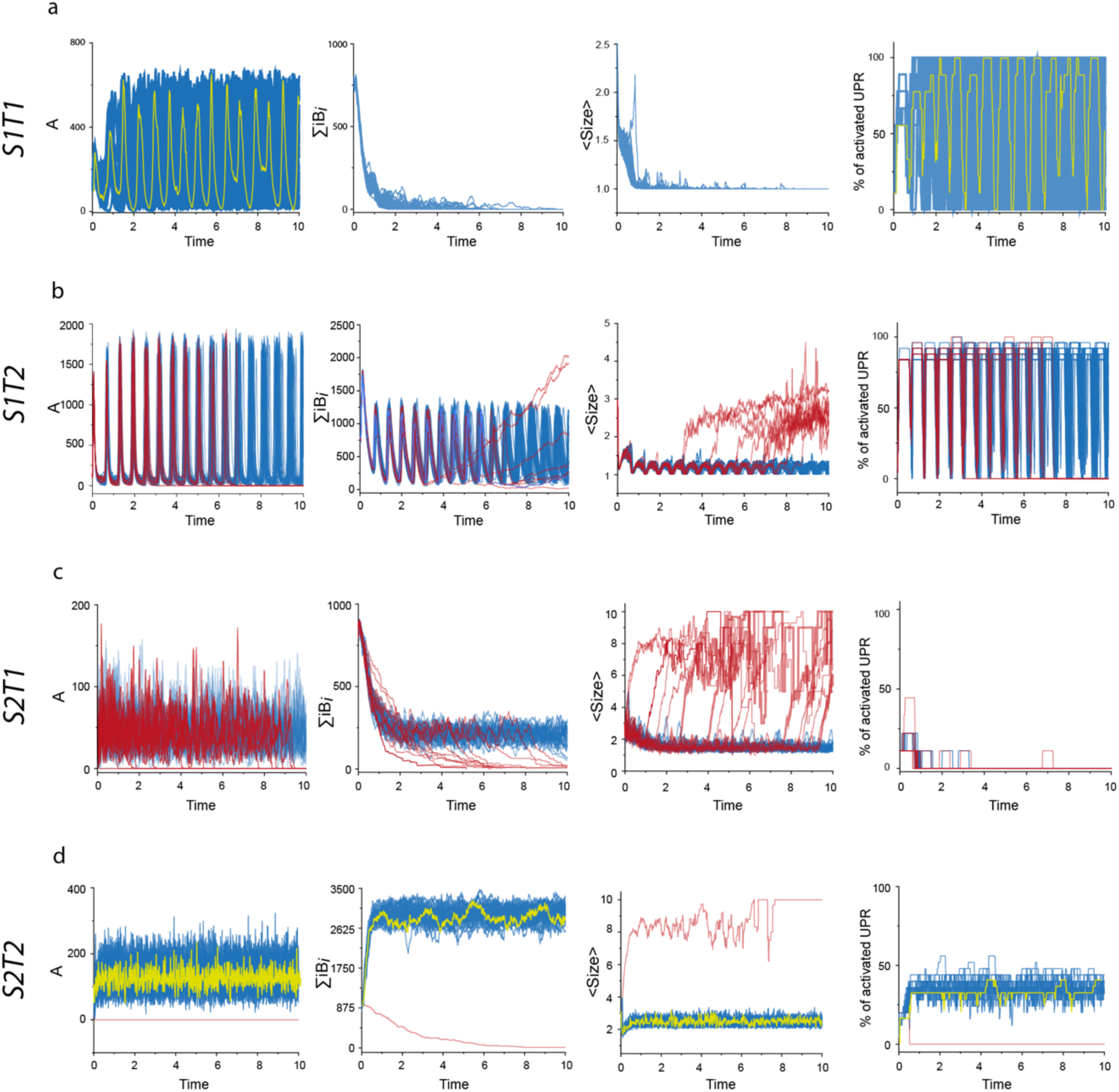
Evolution of two prion strains on two neurons-beds differing in neuron density and UPR activation threshold (see Materials and Methods for more details). These two neurons-beds define tissue 1 and tissue 2. Evolution of different metrics characterizing the process of replication and its sustainability: *A*, ∑ *iB_i_*, average size of assemblies (*<SIZE>*) and percentage of neurons with activated UPR. Rows (a) and (b) correspond to the evolution of strain 1 on tissue 1 (*S1T1*) and tissue 2 (*S1T2*) respectively while (c) and (d) represent the evolution of strain 2 on tissue 1 (*S2T1*) and tissue 2 (*S2T2*) respectively. For each of the four strain/tissue combinations, fifty independent replicates were computed. In all panels, replicates where population *A* was maintained during the simulation timescale are represented in blue while red curves correspond to replicates where *A* was eliminated before the end of the simulation. In rows (a) and (d) where almost all replicates converge, a typical evolution is highlighted in yellow.

**Fig.3.**
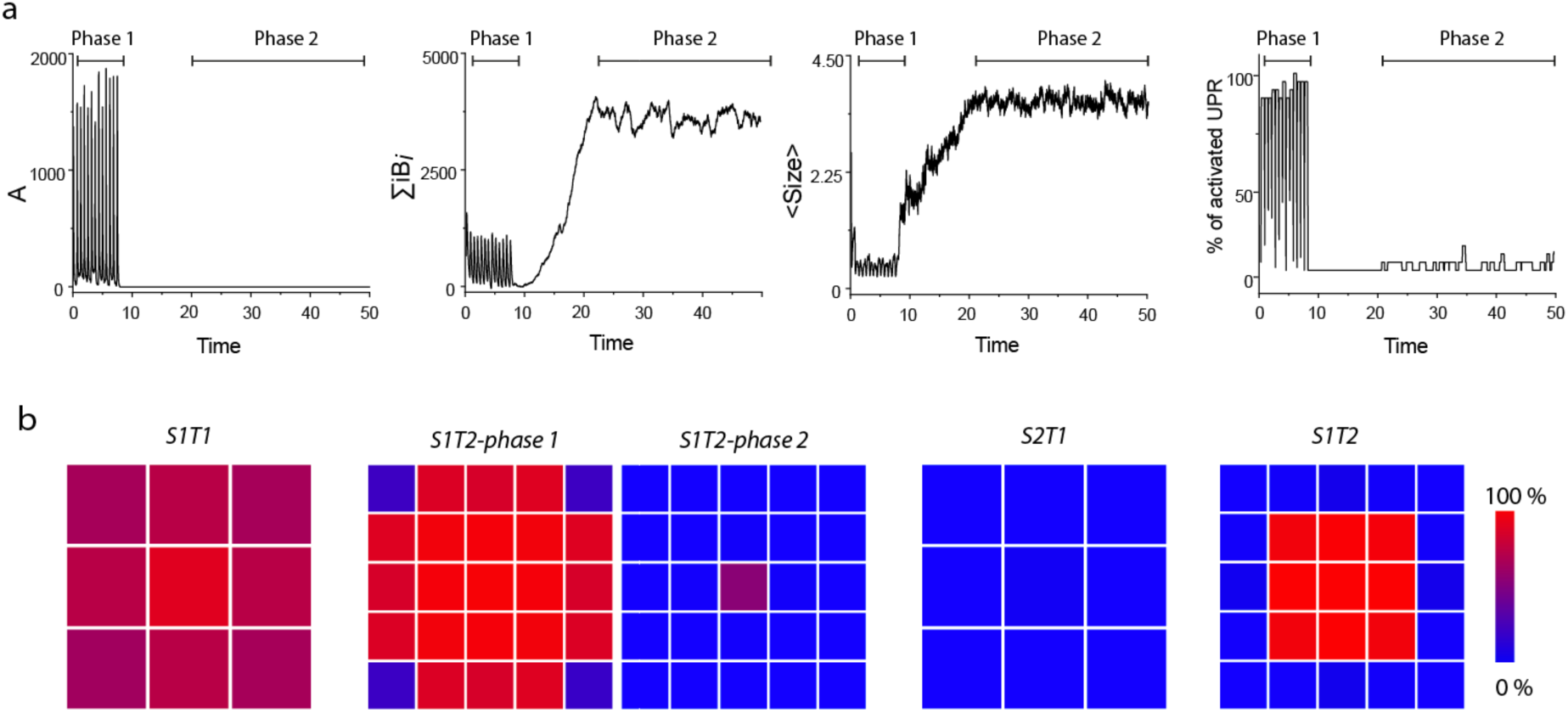
(a) An extension of one replicate of *S1T2* (Fig.2b) with the same metrics highlighting the bifurcation between the two states depending on the elimination of *A*. The two phases are indicated above each panel. (b) Spatial pattern of the average percentage of simulation time spent in UPR activated state for each neuron over fifty independent replicates. (*S1T1*) and (*S2T1*) correspond, respectively, to the evolutions of strain 1 and strain 2 on tissue 1, which is a 3×3 neuron grid. (*S1T2-phase1*/*S1T2-phase2*) and (*S2T2*) correspond, respectively, to the evolutions of strain 1 and strain 2 on tissue 2, which is a 5×5 neuron grid. *S1T2* has two patterns corresponding to the two equilibrium phases previously described. The first one (*S1T2-phase1*) is averaged between the start of the simulation and the moment *A* is eliminated. The second pattern (*S1T2-phase2*) is obtained by averaging the signals from the first UPR activation after *A* has been eliminated to the end of the simulation.

The same strain on tissue 2 (*S1T2*) behaved differently, highlighting the impact of the tissue in the evolution of prion assemblies. While the *B_i_* assemblies were rapidly and systematically eliminated in *S1T1*, *S1T2* presented a transient equilibrium where both its *A* and *B_i_* subpopulations were maintained (Fig.2b and SI3). This state appeared unstable since the proportion of replicates eliminating their *A* subassembly increased with simulation time (SI6). Once the *A* population was eliminated, the quantity and size of *B_i_* assemblies increased to reach what appeared to be a new stable steady-state (Fig.2b and Fig.3a). In the transient state where species *A* and *B_i_* coexisted, an oscillatory out-of-phase behavior based on replication and activation of the UPR was observed for both subpopulations. However, unlike *S1T1*, the UPR of the neurons were not all synchronized (Fig.2b and Fig.3b). Notably, the average activation gradient observed on tissue 1 was much steeper as the UPR of the neurons in the center of the neuron grid occasionally did not deactivate while, in the corner, they were not triggered as much despite being located at the same distance from the edges of the domain (Fig.3b). Once population *A* was eliminated, the UPR signal deteriorated, not triggering until the new equilibrium was reached and then only showcasing impulses from a few neurons in the middle of the neuron bed (Fig.3b).

On tissue 1, the evolution of strain 2 (*S2T1*) was more chaotic than that of *S1T1* (Fig.2c and SI4). The *A* and *B_i_* subassemblies transiently coexisted in roughly constant quantities in the same order of magnitude as *S1T1*. As with *S1T2*, this equilibrium appeared unstable and presented a bifurcation based on the elimination of species *A* (SI6). Once this happened, after a transient increase in average size, the *B_i_* subpopulation was also gradually eliminated leading to an abortive replication in the timescale of the simulations. Interestingly, despite transient replication, the evolution of *S2T1* barely triggered any UPR response from the neurons (Fig.2c and Fig.3b).

The propagation of strain 2 on tissue 2 (*S2T2*) presented a similar pattern of accumulation compared to *S2T1* with roughly constant quantities of *A* and *B_i_* assemblies but with a ratio favoring the *B_i_* subpopulation (Fig.2c and SI5). The average size of the assemblies was also significantly higher. *S2T2* did trigger a UPR response but, unlike *S1T2*, there were no oscillations. In particular, the UPR of the neurons in the center were almost always active while the neurons on the edges barely reacted to the assemblies (Fig.3a). Unlike *S2T1*, this behavior was observed in almost all replicates (49/50).

These observations indicate that the resultant of prion replication within a brain area is influenced by both intrinsic kinetics of the assemblies, specific to a given prion strain, as well as local tissue parameters, specific to a group of cells. Notably, the complex dynamicity between several subpopulations brought out vastly different behaviors depending on the combination of strain and tissue parameters, including selection of one of the subpopulations or transient equilibriums leading to elimination of assemblies. The simulations also showcased vastly different tissue reactions to prion replication: absence of response, synchronized UPR spikes or continuous activation of a specific group. These results can provide an answer to strain phenotypes.

The influence of the initial seeding concentration on the evolution of the system was also investigated. With this in mind, we repeated the 50 replicates of our four tissue-strain combinations with an initial quantity of each assembly increased 100-fold. The only simulations affected by this change in initial quantity were the ones of *S1T1*. For all three other combinations of parameters (*S1T2*, *S2T1* and *S2T2*), the initial condition change did not notably impact the evolution of the system (Fig.4). After short transient phases during which the UPR of the neurons all remained activated, the simulations matched their respective behaviors at nominal initial conditions (Fig.2). Indeed, *S1T2* and *S2T1* presented seemingly unstable states leading to the selection of subpopulation *B_i_* or the elimination of both subassemblies, respectively, while replicates of *S2T2* reached the same equilibrium obtained for nominal conditions (SI6 for comparison between initial seeding conditions). For *S1T1*, the increase in initial quantity caused most replicates (47/50) to eliminate their *A* population, shortly followed by the *B_i_* assemblies. Interestingly, the 3 replicates that showed sustained replication reached the same equilibrium as for nominal initial conditions.

**Fig.4.**
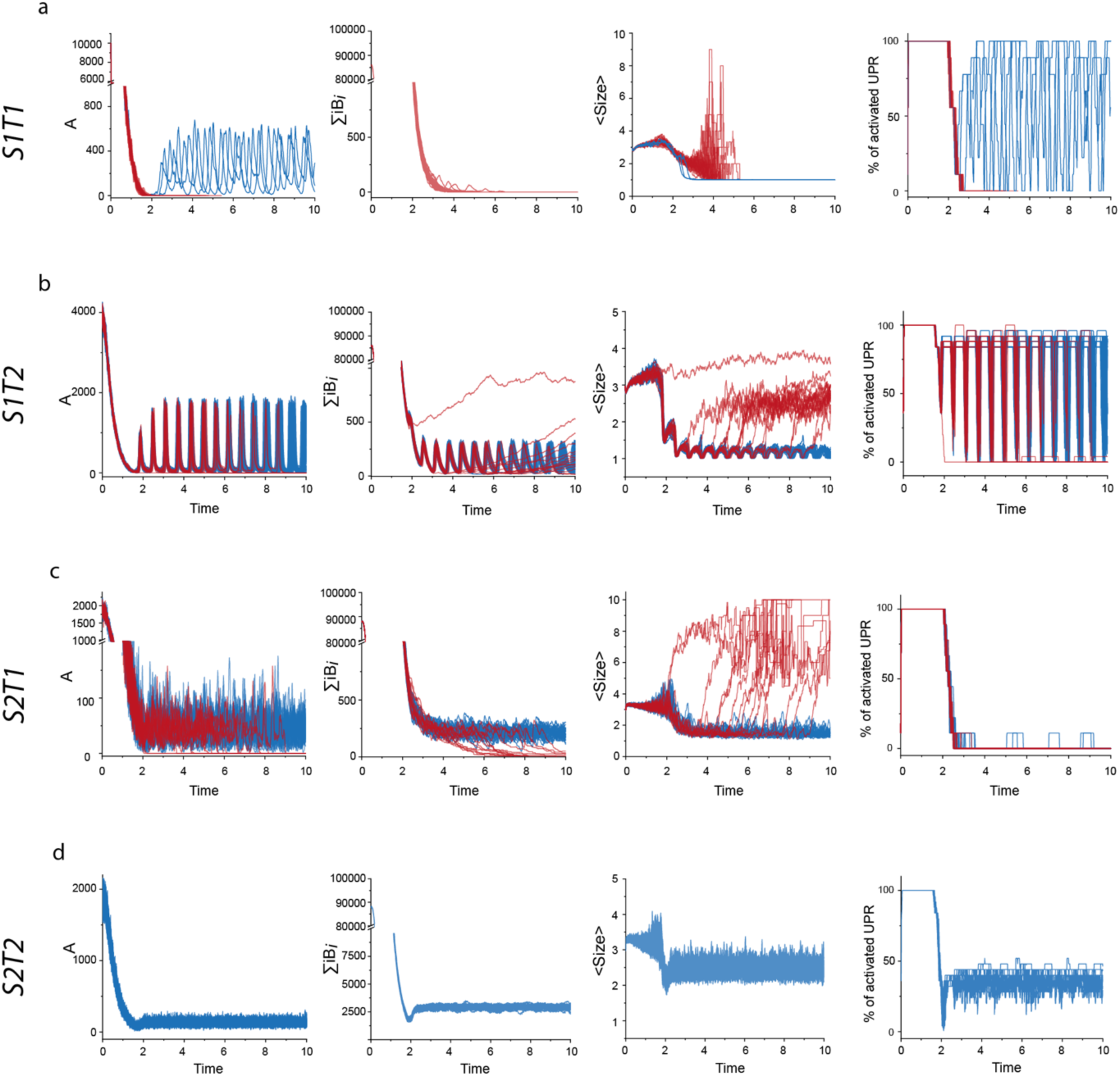
Effect of the initial seed amount on the evolution of strain 1 and strain 2 on tissue 1 and tissue 2. As in Fig.2, the evolutions of *A*, ∑ *iB_i_*, average size of assemblies (*<SIZE>*) and percentage of neurons with activated UPR are represented for 50 independent replicates. Rows (a) and (b) correspond to the evolution of strain 1 on tissue 1 (*S1T2*) and tissue 2 (*S1T2*) respectively while (c) and (d) represent the evolution of strain 2 on tissue 1 (*S2T1*) and tissue 2 (*S2T2*) respectively. Compared to the results on Fig.2, increasing the seed amount does not appear to impact the final equilibriums but, for *T1S1*, most replicates eliminate their *A* subassemblies and fail to reach the outcome previously observed. The comparison between initial seeding conditions can be further analyzed with the help of SI6 showing the percentage of replicates maintaining their *A* subassemblies as a function of simulation time.

While the equilibriums seem governed solely by strain and tissue parameters, the initial quantity of assemblies appears to impact the transient evolution for certain strain-tissue combinations, even leading to the elimination of assemblies for *S1T1* and preventing it from reaching its equilibrium.

### Co-propagation of two strains

As shown in the previous section, the crosstalk between the dynamicity of PrP^Sc^ assemblies and tissue response conditions the sustainability of the replication process and the selection PrP^Sc^ subassemblies.

These observations raise the question of how the tissue response to prion replication could affect the co-propagation of two strains, with possibly different templating activities, as observed in prion field isolates and affected mammalian species ^40,47^.

To address this question, we adapted the previous framework used to compute the evolution of prion assemblies on a homogeneous neuron bed to include two strains. While they are kinetically independent from one another, they both equally contribute to the UPR activation of the neurons during the simulations: the UPR triggers when the sum of assemblies from the two strains exceeds the threshold. We used the simulation parameters from previous sections defining tissue 1 and tissue 2 (Fig.1b) and computed the co-propagation of strain 1 and strain 2 from previous simulations with various templating rate combinations. While the kinetic parameters governing the dynamicity of the strains were kept invariant, the previous templating values are labeled as nominal (denoted as *N*) and we introduce lower (*L*) and higher (*H*) templating rates. Fig.5a summarizes all the different templating combinations computed on both tissue 1 and tissue 2. Moreover, to determine how strain 1 and strain 2 interfere during their co-propagation, we systematically compared the evolution of each strain taken individually to the co-propagation condition.

**Fig.5.**
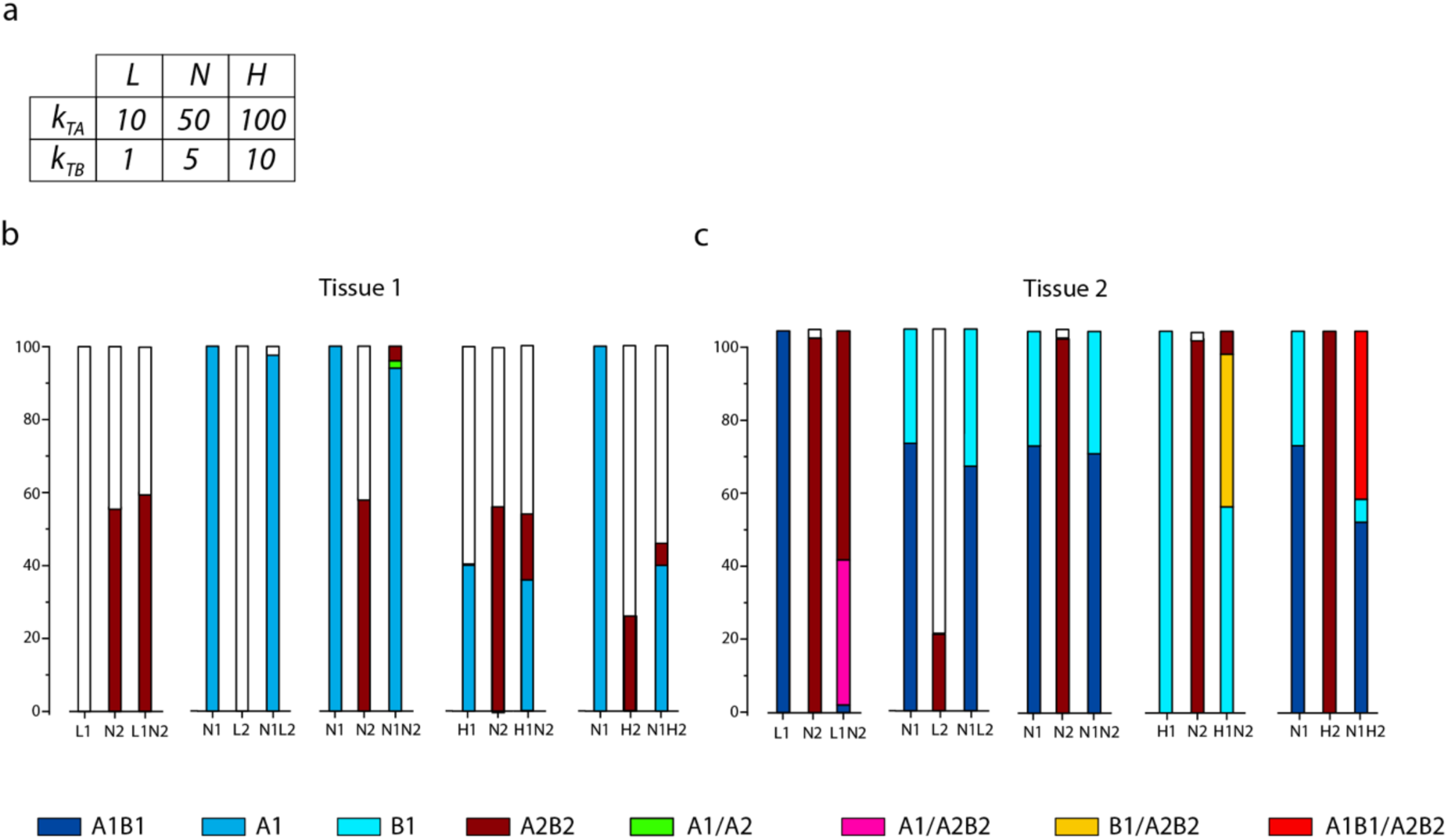
Effect of the replication rate on the evolution and co-propagation of strains on tissues 1 and 2. (a) By taking as a reference the templating parameters of strains 1 and 2 described in Fig.1b (named *N1* and *N2* respectively), we either doubled the replication rates for both *A* and *B_i_* (*H1* and *H2* for strain 1 and strain 2 respectively) or divided them by 5 (*L1* and *L2* for strain 1 and strain 2 respectively) to get the high and low templating configurations for both strains. The evolutions of five different strain combinations were then computed and summarized on graph (b) for simulations on tissue 1 and graph (c) for tissue 2. The co-propagation of a strain combination is grouped with the two strains individually for ease of comparison. The bars represent the outcomes of 50 independent replicates, white means no assembly sustainably replicated within the simulation timeframe.

On tissue 1, as shown in the previous section, the replication of strain 1 and strain 2 with nominal templating rates (denoted as *N1* and *N2*) led to sustainable replication in the timescale of the simulations in respectively 100% and 58% of the replicates. For strain 1, *B_i_* assemblies were eliminated while strain 2 maintained both its subpopulations in the sustainable replicates (Fig.2a&c and Fig.5a). Lowering the templating rate (denoted as *L1* and *L2* for strain 1 and strain 2 respectively) led to an unsustainable replication for both strains in 100% of the replicates (Fig.5b). In copropagation conditions with the other strain at nominal rate (*L1N2* and *N1L2*), the resultant was very similar to the behavior of the nominal strain by itself. Indeed, for *L1N2*, 60% of the replicates led to the same apparent outcome that was observed in 58% of the replicates with *N2* alone. For *N1L2*, almost all replicates (98%) ended with the selection of subpopulation *A* from strain 1 similarly to the results of *N1* alone. The *L1N2* and *N1L2* conditions illustrate a case where the tissue systematically selects the better replicant with little interference between the two strains.

With both templating rates set to nominal (*N1N2*) on tissue 1, the *A* subpopulation of strain 1 was predominantly selected in 94% of the replicates against 4% for strain 2. Interestingly, 2% of the simulations resulted in a sustained copropagation of the two strains. Despite both strains sharing the same templating rates, the quasi-total selection of strain 1 in this copropagating configuration highlights the contribution of kinetic parameters governing the intrinsic dynamicity of PrP^Sc^ assemblies in the co-propagation of two strains.

Increasing the templating rate of the two strains (respectively denoted as *H1* and *H2*) unexpectedly led to a decrease in sustainable replication on tissue 1 for both strains compared to their nominal evolution, all while selecting the same subpopulations as reported in Fig.5b (40% in *H1* and 26% in *H2*). In copropagating condition *H1N2*, simulations showed the selection of mostly strain 1 (36%) with strain 2 accounting for 18% of replicates. While the number of replicates favoring strain 1 was similar to the evolution of strain 1 alone in the *H1* configuration, the number of sustained replicates of strain 2 dropped from 58% in *N2* to 18% in the copropagating condition *H1N2*. Here, despite strain 1 with a high templating rate (*H1*) being less sustainable than nominal strain 2 (*N2*) on their own, in copropagating conditions, *H1* was selected more at the cost of *N2*. Finally, while *N1* resulted in 100% maintained evolutions on its own, computing its evolution with *H2* (26% on its own) resulted in less than half of the replicates showcasing sustained replication: *N1H2* configuration ended in 40% strain 1 selection and 6% strain 2 selection. In this example, despite not interacting through any kinetical pathway, both strains exerted negative feedback on the replication of the other as each strain ended up with fewer sustained evolutions combined than on their own.

On tissue 2 (Fig.5c), at nominal templating rate (*N1*), strain 1 was maintained in 100% of the replicates but with two different outcomes observed in the timescale of the simulations: in 74% or replicates, *A* and *B_i_* subassemblies were maintained while in the remaining 26%, only subpopulation *B_i_* remained. The replication of strain 2 at nominal configuration (*N2*) on tissue 2 resulted in 98% of the replicates maintain both subassemblies. As on tissue 1, decreasing the templating rate of strain 2 on tissue 2 (*L2*) significantly decreased the number of sustainable replicates (20%) compared to the *N2* configuration. In the low configuration, strain 1 (*L1*) was still maintained in 100% of the simulations, but unlike its evolution in the *N1* configuration, *A* and *B_i_* were persistent in every simulation. Copropagating strain 1 and strain 2 with respectively low and nominal templating rates (*L1N2*) on tissue 2 resulted in a different outcome compared to tissue 1. Notably, even though strain 1 was eliminated in 60% of the replicates, a sustained co-propagation of strain 1 and strain 2 was also observed in 38% of the replicates despite both strains having different templating activities. Additionally, in the replicates that maintained both strains, the equilibrium of strain 1 was shifted, only maintaining its *A* subassembly in the presence of both *A* and *B_i_* subpopulations from strain 2. Here, not only was the number of sustained replicates impacted but also the balance of subpopulations of strain 1, even eliminating one of them. This is another example of strains influencing each other while copropagating despite not being kinetically linked. The copropagating condition *N1L2* on tissue 2 behaved similarly as on tissue 1: strain 1 dominated with little interference from strain 2, even replicating a distribution of outcomes alike nominal strain 1 on its own (*N1*). This observation was also true in the *N1N2* configuration despite *N2* showing sustainable evolution in 98% of replicates on its own.

On its own, when its templating rate was increased, strain 2 (*H2*) showed little difference from its nominal behavior (*N2*) on tissue 2 as its *A* and *B_i_* were maintained in all replicates (Fig.5c). While strain 1 was also maintained in 100% of the simulations at high templating rate on tissue 2 (*H1*), only its *B_i_* subassemblies sustainably replicated unlike in the *N1* and *L1* configurations. In both copropagating configurations including a highly replicative strain (*H1N2* and *N1H2*), sustainable co-propagation of the two strains was observed in quasi-similar proportions. In the *H1N2* condition, copropagating ended in a product of the two strains taken individually: 54% of strain 1 *B_i_* subassemblies selection (like *H1*), 6% of strain 2 with both subpopulations (like *N2*) and 40% of sustained co-propagation with *B_i_* subassemblies from strain 1 and both species from strain 2. In the *N1H2* configuration however, even with strain 2 having a higher templating rate, it was not selected over strain 1 in any replicate. Interestingly, despite strain 1 having 2 outcomes on its own (conservation of both subassemblies or elimination of *A* in *N1*), in all of the 48% replicates that featured sustained co-propagation, both subassemblies of strain 1 were maintained.

Throughout the simulations of the different tissue-templating combinations, we noticed a specific role of the tissue. In the absence of kinetic interaction between the two strains, the tissue mediated their linkage as they equally contribute to the UPR of the neurons. This resulted in the strains being able to influence one another and even led to dominance negative interferences (*N1H2* configuration on tissue 1) or copropagation with the selection of specific subassemblies. The impact of tissue parameters in the co-propagation of two strains was also showcased here as tissue 2 appeared to allow coexistence of strain 1 and strain 2 while tissue 1 did not, even failing to maintain highly replicative strains. Additionally, because of the intrinsic dynamicity of the strains, selection of the worse replicator or sustained co-propagation of two strains with different templating activities was observed.

### Contribution of the connectome to the spreading

While we previously focused on studying the contribution of a modeled brain area to the replication and selection of PrP^Sc^ assemblies, we then explored the influence of the axonal neural network to the dissemination of the prion assemblies for three distinct strains (Fig.1b). While the three strains share the same templating activities, due to their intrinsic dynamicity, the balance between subpopulations is gradually more in favor of *B_i_* from strain 1 to strain 3. Instead of a homogeneous neuron bed, two clusters of neurons are placed on opposite corners of the domain, linked unilaterally either by anterograde or retrogrades axons. Additionally, the number and size of voxels in the square-shaped domain are increased to emulate longer distances. The replication around neurons (somas and axons) is regulated by the UPR triggered once the concentration of PrP^Sc^ reaches a threshold in the vicinity of the associated soma (see Materials and Methods and Fig.1d for more detail).

The replication of three different strains was initiated at one cluster and the anterograde or retrograde dissemination towards the other was monitored using three metrics: the time of arrival at the opposing cluster (*t_arriv_*), the average size (*<Size>*) and quantity of assemblies (*A* + ∑ *iB_i_*) at *t_arriv_* (Fig.6).

**Fig.6.**
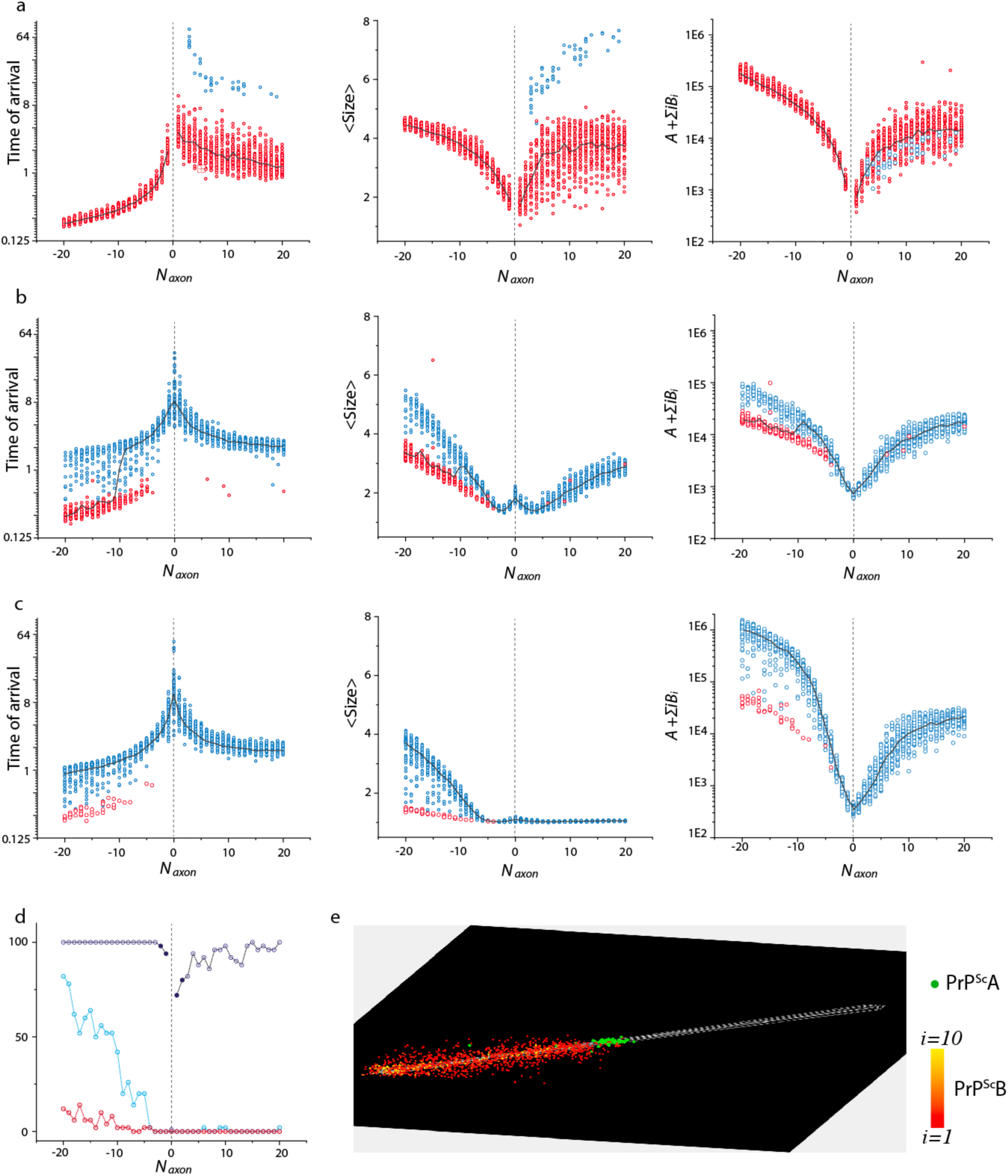
Influence of the number and orientation of axonal connections on the spreading process between two unilaterally linked neuron clusters for three distinct prion strains (see Fig.1b for strain parameters). For all graphs, the x-axis accounts for the number of axons (*N_axon_*) linking the clusters, negative values correspond to retrograde propagation (seeding at the receiving neurons) and positive ones to anterograde (seeding at the emitting neurons). For each connectivity value and each strain, fifty replicates were computed. Their output is represented by dots in the scatter plots of row (a) for strain 1, row (b) for strain 2 and row (c) for strain 3. We studied the influence of connectivity on three different metrics: the time of arrival to the second cluster as well as the average size (<SIZE>) and quantity of assemblies (*A* + ∑ *iB_i_*) at the time of arrival. Blue dots correspond to simulations which eliminated their *A* subassemblies before reaching the second neuron group, while red ones maintained them. The black line is the median value of the replicates for each metric. (d) The proportion of replicates maintaining their *A* subassemblies during the simulation timescale as a function of connectivity (*N_axon_*) for all three strains highlights the impact of axons and UPR response on the sustainable replication of subpopulations. (e) Typical frame taken from a video of the retrograde dissemination process (see SI7 for full video). Seeding was done at the receiving neurons on the left, *A* assemblies are represented in green while *B_i_* assemblies are colored from red to yellow depending on size. This shows that the dissemination was guided by axons and facilitated by the *A* subpopulation being more replicative and diffusive. In this configuration, the system self-organized with a front of *A* followed by the *B_i_* subpopulation, with larger assemblies located closer to the place of inoculation.

Overall axons appeared to reduce the time of arrival for both anterograde and retrograde spreading pathways. In particular, compared to dissemination in the absence of axons, anterograde or retrograde connections significantly decreased *t_arriv_* for the three strains (Fig.6a-c); this was especially true for strain 1 which failed to reach the opposing cluster without axons (Fig.6a). This indicates that the connectome induced a super-diffusion phenomenon ^48^ due to replication along the axon’s path, both in anterograde and retrograde progression. It is also worth noting that for all three strains, for a given number of axons, retrograde spreading was faster than anterograde on average. This is because the UPR was not triggered until the second cluster was reached during retrograde dissemination, meaning that the replication along the axons was unregulated during the simulation.

By first analyzing the strain effect on the anterograde dissemination pathway, we observed that a few replicates of strain 1 eliminated their *A* subpopulation by the time they reached the second group of neurons (Fig.6a). These replicates presented a significantly higher *t_arriv_* and average size compared to the ones where *A* was maintained but they did not stand out when it came to quantity of assemblies. As anterograde connectivity increased, the number of strain 1 replicates which maintained their *A* subpopulation during the dissemination increased (Fig.6d). Conversely, strain 2 only maintained *A* during anterograde spreading in a few replicates without any apparent sign of increase with connectivity (Fig.6d). As was the case with strain 1, these replicates also presented lower *t_arriv_* than those where *A* was eliminated (Fig.6b) but, unlike strain 1, they did not correspond to a lower average size of the assemblies or a lower quantity of assemblies. When it came to strain 3 (Fig.6c), the anterograde spreading process systematically eliminated the *A* subpopulation before reaching the second cluster (Fig.6d). While the evolutions of time of arrival and quantity of assemblies shared similarities with the majority of the other 2 strains, the average size of strain 3 assemblies in anterograde spreading did not appear to depend much on the number of axons, always favoring small *B_i_* assemblies.

In retrograde dissemination, strain 1 replicates showed a uniform behavior as all of them maintain their *A* subpopulation (Fig.6d). The retrograde spreading was faster and accumulated more assemblies than the anterograde while maintaining a similar size distribution (Fig.6a). Strains 2 and 3 however presented a dual behavior based on the elimination of *A* in retrograde spreading (Fig.6b&c). At low retrograde connectivity, all replicates of strains 2 and 3 eliminated *A* during their dissemination to the second cluster but the proportion of replicates which maintained *A* increased with the number of axons, faster for strain 2 compared to strain 3 (Fig.6d). For both strains, when *A* was maintained, the speed of dissemination notably increased, reaching that of strain 1 while the average size and number of assemblies were lower (Fig.6b&c). Out of the three strains, strain 3 showed the biggest differences based on spreading pathway, accumulating more and much larger assemblies in the retrograde direction.

The existence of two distinct regimes raises questions about the spatial distribution and evolution of *A* and *B_i_* subpopulations in the regime where *A* is maintained. As shown in the time-lapse spatiotemporal retrograde spreading of strain 1, a dissemination front of *A* assemblies preceded *B_i_* subassemblies (Fig.6e, for the full movie see SI7). This faster spreading of population *A* appeared to facilitate the following dissemination of *B_i_*, leaving behind a trail of condensation with bigger and more numerous assemblies located near the starting point.

Taken together, these observations suggest that the density of the connectome and the spreading pathway (anterograde or retrograde) interfere with the intrinsic strain properties to determine the speed of dissemination as well as the nature, size and number of assemblies. Once again, we highlighted a key role of the *A* subassemblies as their retention, based on connectivity and strain kinetics, greatly impacts the dissemination process.

## Discussion

The link between replication and dissemination of prion assemblies through the brain tissue is pivotal to our understanding of prion pathologies. However, this coupling is poorly understood and PrP^Sc^ deposition pattern, unlike other proteinopathies. While the connectome is a key factor in the pathogenesis of Alzheimer’s and Parkinson’s diseases ^15,16^, its contribution to PrP^Sc^ deposition patterns is unclear ^17^ as the evolution of prion diseases and dissemination of the assemblies seem more complex and strain dependent.

Prior to this work, several approaches had been developed to explore the relation between prion replication and tissue spreading to no avail ^55^. These often-featured nucleation-elongation kinetics, imposed spreading through the structural connectome or lacked any form of tissue response to prion replication. Unlike the previous studies, our model is based on recent developments in the mechanisms of prion replication, featuring multiple subpopulations engaging in autocatalytic processes, and includes an albeit simplistic tissue response to prion propagation in the form of a negative feedback impacting PrP^C^ production via the UPR.

By biochemically formalizing prion strains as the convolution of intrinsic dynamicity of PrP^Sc^ assemblies and their replicative properties, we demonstrated that PrP^Sc^ replication, accumulation and neuronal response within a brain area depend non-linearly on both strain and tissue properties. This crosstalk between the dynamicity of PrP^Sc^ assemblies and the tissue response conditions the sustainability of the replication process as well as the co-propagation of different PrP^Sc^ subpopulations or distinct strains with different templating activities in coinfection conditions. We also present in this work a valid alternative to step-by-step spreading through the structural connectome, in the form extra-cellular diffusion with replication in the vicinity of neurons.

### The tissue response is an integral part of the prion replication process

The modification of the structural and functional neural networks, neural death, astrocytosis, UPR activation and metabolic response are all part of the complex and intricate tissue response to prion replication ^56^. Despite this complexity, we chose to only consider the UPR in this work to simplify the model as it is one of the earliest responses to prion propagation and its mechanisms have been extensively studied ^33–37^. It has been reported that accumulation of PrP^Sc^ locally triggers the UPR of neurons which results in the downregulation of proteins transiting through the endoplasmic reticulum, including PrP^C^ ^37,57^. This negative feedback of PrP^Sc^ on PrP^C^ production can interfere with prion propagation by introducing non-linearity in the replication process ^52^. Considering other types of tissue responses or a combination of them could increase the non-linearity of the response to prion replication therefore further complexifying the system and allowing for the emergence of new behaviors.

The tissue response to PrP^Sc^ accumulation is also known to be cell-specific. Different types of neurons may exhibit different thresholds of UPR activation, with downstream effects varying depending on cellular context ^58,59^. This makes the UPR a suitable candidate for a local tissue parameter differentiating between brain areas. In our model, the interruption of the templating due to the accumulation of assemblies can also be interpreted as PrP^C^ becoming the limiting factor in the replication process once the concentration of PrP^Sc^ reaches a certain threshold. This would also qualify as a local tissue parameter as the latency in PrP^C^ production could fluctuate between cell types and cause similar non-linearities. As typically shown in the evolution of *S1T1* and *S1T2*, the oscillatory behavior of the replication is a direct consequence of the negative feedback of UPR on PrP^C^ production. However, the absence of periodic patterns in the accumulation of *S2T1* and *S2T2* shows that the UPR is not the sole factor. In our simulations, the oscillations in PrP^Sc^ quantity systematically occur in conditions where the proportion of *A* subassemblies is important, such as *S1T1* or *S1T2*. In those situations, the tissue response to the accumulation consists of synchronized UPR firings of the neurons. Conversely, simulations with higher proportions of *B_i_* subassemblies do not exhibit oscillatory behaviors of PrP^Sc^ quantity or UPR signals, even showcasing a temporary state where the replication does not elicit any tissue response in the case of *S2T1*. While the high replicability and diffusivity of *A* induces synchronization between all neurons, the inertia caused by the lower diffusivity and condensing of *B_i_* decreases the coupling distance, resulting in the continuous stimulation of neurons in the middle of the grid, thereby causing the emergence of spatial patterns. As the balance between *A* and *B_i_* subpopulations depends partially on the strain, this indicates that PrP^Sc^ accumulation results from an interplay of the tissue response and intrinsic dynamic of strain replication.

As some propagation simulations of *S2T1* or *S1T2* showed, we found instances of stable or transient unstable replication during which the UPR is barely triggered or not triggered at all. Even if the tissue response is extremely simplified in this model, being restricted to the UPR activation, this particular behavior tends to indicate that silent replication escaping the tissue response could occur for some combinations of tissue parameters and strain dynamicity.

### Entanglement between PrP^Sc^ dynamicity and tissue response governs tissue tropisms, selection of subassemblies and co-propagation

It is commonly thought that the initiation and sustainability of prion replication in a given tissue is governed by the compatibility between PrP^Sc^ assemblies and the local conformome of PrP^C^ or local cofactors thermodynamically influencing the homotypic replication process. Here, our simulations revealed that introducing non-linearity in the form of a tissue response to prion replication modulating PrP^C^ expression could impact the sustainability of the replication process and the selection of PrP^Sc^ subassemblies. As shown in Fig.2 and Fig.3, with limited strain-tissue parameter combinations, we reached a wide variety of outcomes such as the selection of a specific PrP^Sc^ subpopulation depending on tissue (*S1T1* vs *S1T2*) or transient unstable replication (*S2T1*). We also showed that for some strain-tissue combinations, increasing the replication rate (Fig.4) or initial seed concentration can negatively impact the sustainability of the replication process, as highlighted by the behavior of *S1T1* (Fig.2 and Fig.3). These observations indicate that the resonance between a non-linear tissue response and the inherent dynamic nature of prion assemblies could explain strain-specific tissue tropisms and provide a viable alternative to the local conformome hypothesis.

Our simulations also revealed the existence of transient replication regimes. The evolutions of *S1T2* and *T1S2* present initial states where both subpopulations are transiently maintained. However, both these equilibria appear unstable as they eventually lead to the elimination of *A* subassemblies as simulation time increases (SI6). Due to the highly dynamic nature of the kinetic scheme, the elimination of one of the subpopulations results in a drastic change in behaviors. For *S2T1*, the loss of subpopulation *A* is shortly followed by that of *B_i_*, resulting in abortive replication, while *S1T2* reaches a new equilibrium composed solely of *B_i_* subassemblies. Despite similarities, the two transient replication regimes elicit radically different responses from the neurons: *S1T2* presents periodic firing of the UPR of most neurons while *S2T1* provokes no response. This shows that transient replication relies on both strain dynamicity and the tissue-modulated replication, highlighting once again this interplay as a key factor in prion propagation. These transient replication regimes could explain experimentally observed non-adaptive prion amplification ^60^.

The non-linearity introduced in our model by the tissue response also plays a key role in the co-propagation of two prion strains within the same brain area. We showed that the copropagation of two strains can alter their respective evolution or sustainable replication, giving rise to negative dominance interferences ^61,62^. As we made the strains kinetically independent in our model, this interference is a direct result of the competition for the same substrate and equal contribution to the UPR of the neurons highlighting the influence of the tissue response in the strain selection process. The tissue also appears to play a crucial role in the coexistence of two strains. As shown in Fig.5b and Fig.5c, tissue 1 does not allow the sustained co-propagation of any combination of our two modeled strains while tissue 2 does. This type of behavior is observed experimentally in prion field isolates where two strains can coexist within the brain while other organs such as the spleen only maintain one of them ^63^. Combined with the non-linear tissue response, the intrinsic dynamicity of prion assemblies, reflected in the kinetic scheme comprising several autocatalytic reactions between different subpopulations, allows for possible best replicator elimination or sustained co-propagation of strains with different templating activities. This provides answers to strain selection, co-propagation and negative dominance effects that are not based on hypothetical cofactors or structural compatibility between PrP^Sc^ and tissue-specific PrP^C^ conformers (thermodynamic considerations) but on kinetic considerations and complex system responses ^64^.

### Contribution of the connectome to the process of spreading

One of the main questions in the field of prions is how the connectome contributes to the spreading and neuro-invasion process. Experimental observations indubitably highlighted the impact of both peripheral anterograde and retrograde connections ^65,66^ while excluding axonal cytoskeleton transport ^66^. In our simplified configuration featuring two groups of neurons linked unilaterally, we showed that axons facilitate prion dissemination and promote the replication of assemblies, increasing both quantity and average size. Unlike previous works imposing axonal transport, in our model the assemblies are only guided by the connectome as they disseminate according to Brownian motion and can replicate in the vicinity of axons. In this context, despite different tissue configurations and kinetic parameters, the sustainability of *A* subassemblies once again emerged as a key factor in the spreading process, acting as a switch between two distinct regimes notably differing in propagation speed. If subpopulation *A* is maintained during propagation, for a given strain and connectivity, the dissemination speed increases greatly while the quantity and size of assemblies at the time of arrival is lowered. This role of facilitator is due to its high diffusivity and replicability combined with its special role in the highly catalytic kinetic scheme. This is reflected in the spatial organization of assemblies that emerges during the dissemination process: a front of *A* followed by a gradient of size of *B_i_* subpopulation, with the smaller *B_i_* assemblies following due to *B_1_* being a predator of *A* (SI7). This is why the retention of *A* by the time the second neuron group is reached depends on the intrinsic dynamicity of the strains as well as the connectivity, with axons contributing to the sustainability of *A*, especially in retrograde spreading. Due to the progressive shift in behaviors occurring as retrograde connectivity increases, once again we highlighted a resonance between tissue and strain parameters resulting in non-linearities when it comes to propagation speed and accumulation. Simulations also revealed a potential role of the connectome as a filter in the spreading process, selectively determining the types of assemblies to propagate. This selection process could determine which assemblies initiate replication in the next region.

Additionally, one of the main conclusions emerging from this study is the difference between retrograde and anterograde disseminations. In all our simulations, strains propagate faster in the retrograde direction, this is exacerbated at high connectivity due to the previously highlighted shift in regimes mainly occurring in the retrograde direction. Strains also appear to react differently to the direction, for instance, the quantity of assemblies accumulated for strain 2 seems relatively independent of direction while strain 3 accumulates significantly more. The difference in speed depending on directions is not experimentally documented and could be a direct consequence of our modeling of the UPR. In our retrograde propagation model, the UPR associated with the axons is not triggered until the second group of neurons is reached (see Materials and methods), resulting in an unregulated replication greatly accelerating the dissemination process. Conversely, in the anterograde direction, the presence of aggregates around the somas of the first group of neurons immediately prevents replication on the axons. While it is possible that the assemblies going up the axons would not elicit any tissue response, the replication process would be expected to decline due to the high number of assemblies causing the production of PrP^C^ to become the limiting factor. This would not impact the dissemination front thus we can expect the speed to be affected less than the quantity or the size of assemblies.

## Conclusions

In the present work, we showed that a combination of non-linearities imposed by tissue response and highly dynamic prion kinetics can make prion propagation in the brain tissue a complex system. This complexity resulting from the entanglement between negative feedback on PrP^C^ expression and multiple catalytical processes between different PrP^Sc^ subpopulations promotes the emergence of unpredictable behaviors such as specific subpopulation selection, interference between two strains with negative dominance as well as abortive, transient or silent replication. This research provides an alternative to a hypothetical variable compatibility between strains and local conformers of PrP^C^ to explain strain specific phenotypes and tissue tropisms which could not be resolved otherwise with the canonical linear ways to consider prion replication (end-elongation or nucleation-elongation processes).

The simulations further clarify the connectome’s involvement in the dissemination process, showing that it operates not through axonal transport but rather through extra-cellular diffusion and replication near neurons. Its interaction with highly dynamic prion kinetics once again leads to non-linear behaviors in dissemination speed and accumulation as well as spatial organization of the assemblies.

## Acknowledgments

This work was supported by grants from ANR (PrionDif, ANR-21-CE15-0011-01), the European Research Council (ERC Starting Grant SKIPPERAD, number 306321), the Ile-de-France region (DIM MALINF) and Metaprogram DigitBio (PrionDiff).

## Author contributions

HR, BF and LPM designed research; BF and HR performed research and analyzed data; HR, BF, AI, VB, DM, PS performed experimental researches and confronted simulations to experimental observations. HR and BF wrote the paper.

## Supplementary information legends

**SI1:** Simulations codes

All codes used in this work can be uploaded through the following link: https://doi.org/10.5281/zenodo.11093945

**SI2, SI3, SI4 & SI5:** Video of strain propagation on a homogeneous neuron bed. Three metrics (*A*, ∑ *iB_i_*, UPR) are graphed on top with animated vertical time-lines indicating at each instant the corresponding moment represented in the video under. Neurons are displayed as crosses, colored in white when their UPR is inactive and pink otherwise. The assemblies are colored dots, green ones are *A* and *B_i_* range from red to yellow depending on size.

**SI6:**
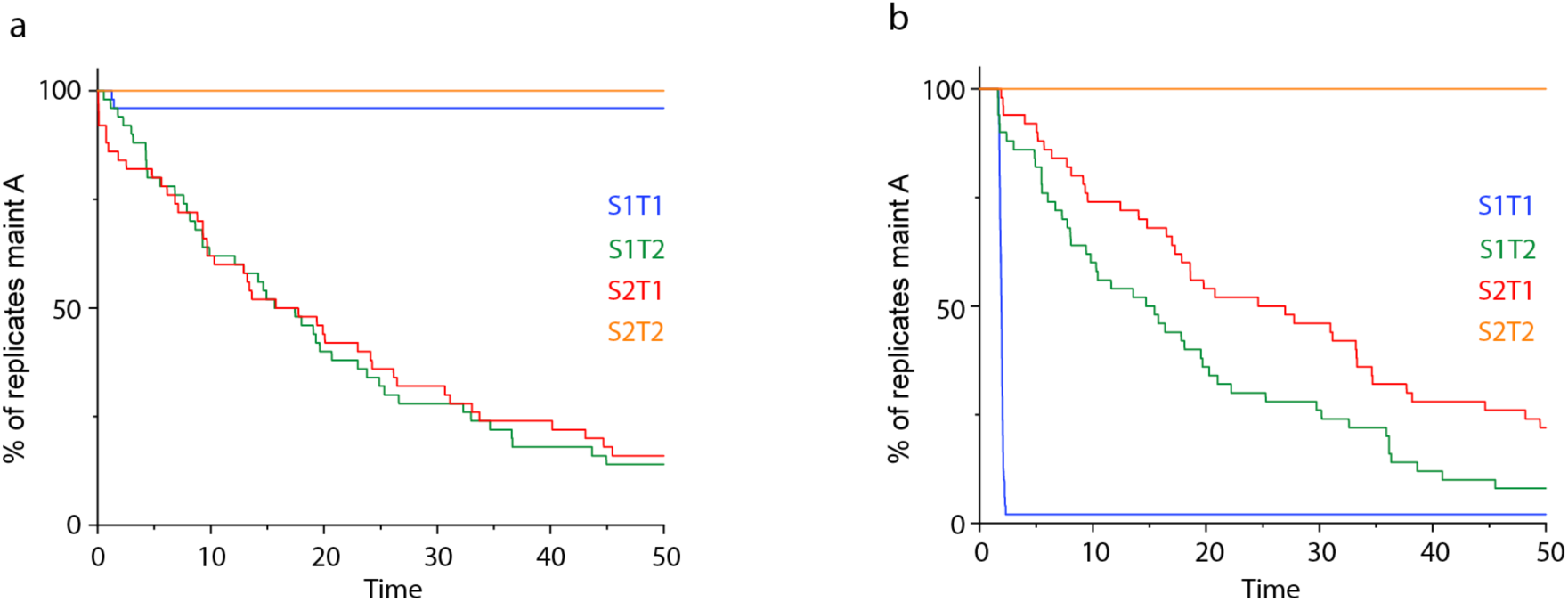
Percentage of replicates maintaining their *A* subassemblies across 50 independent replicates over time for each strain-tissue combination. Graph (a) corresponds to the nominal initial conditions and graph (b) the increased seeding. Both graphs highlight the transient replication state observed with the evolutions of *S1T2* and *S2T1*, as replicates progressively eliminate subpopulation *A* as simulation time increases. The only notable difference between the two graphs is in the evolution of *S1T1*, where the high initial conditions caused most replicates to eliminate their *A* subassemblies early in the simulations. This shows that initial conditions can impact the evolution of certain strain-tissue combinations.

**SI7:** Video of the retrograde dissemination process between two neuron clusters. The two neuron clusters were located on opposite corners of the domain and unilaterally linked by axons (see Materials and Methods for more details). Seeding of the assemblies was done at the receiving neuron cluster. Neurons are represented as white crosses, axons as white lines. *A* assemblies are represented as green dots while *B_i_* are colored using a red-yellow gradient to account for size, red corresponds to small *B_i_* assemblies and yellow big ones. This video highlights the impact of replication on the dissemination process as templating of the assemblies occurred only in the vicinity of the axons. The role of both subassemblies was also brought out as the system spatially self-organized with a front of *A* assemblies, more replicative and diffusive, followed by a gradient of *B_i_* assemblies, with larger objects located closer to the inoculation site.

